# The neutrophil antimicrobial peptide cathelicidin promotes Th17 differentiation

**DOI:** 10.1101/2020.04.10.035543

**Authors:** Danielle Minns, Katie J Smith, Virginia Alessandrini, Gareth Hardisty, Lauren Melrose, Lucy Jackson-Jones, Andrew S. MacDonald, Donald J Davidson, Emily Gwyer Findlay

**Affiliations:** Centre for Inflammation Research, University of Edinburgh; 47 Little France Crescent, Edinburgh, EH16 4TJ; Division of Biomedical and Life Sciences, Lancaster University, Lancaster, UK; Lydia Becker Institute of Immunology and Inflammation, University of Manchester, Oxford Road, Manchester UK

## Abstract

The host defence peptide cathelicidin (LL-37 in humans, mCRAMP in mice) is released from neutrophils by de-granulation, NETosis and necrotic cell death; it has potent antibacterial, antiviral and antifungal activity as well as being a powerful immunomodulator. It is released in proximity to CD4^+^ T cells during inflammatory and infectious disease but its impact on T cell phenotype is scarcely understood. Here we demonstrate that cathelicidin is a powerful Th17 potentiating factor which increases expression of the aryl hydrocarbon receptor (AHR) and the RORγt transcription factor, in a TGF-β1-dependent manner. We show that cathelicidin induces IL-17F production in particular, and that its induction of IL-17A+F+ double producing cells is dependent on AHR while its induction of IL-17F single producing cells is not. In the presence of TGF-β1, cathelicidin profoundly suppressed IL-2 and down-regulated T-bet, specifically directing T cells away from Th1 and into a Th17 phenotype. Strikingly, Th17, but not Th1 cells were protected from apoptotic death by cathelicidin, in the first example of a neutrophil-released mediator inducing survival of a T cell subset. Finally, we show that cathelicidin is released by neutrophils in mouse lymph nodes following inoculation of heat-killed *Salmonella typhimurium* and that cathelicidin-deficient mice have suppressed Th17 responses during inflammation, but not at steady state. We propose that the release of cathelicidin by neutrophils is required for maximal Th17 differentiation and IL-17 production by CD4^+^ T cells, and that this is one method by which early neutrophilia directs subsequent adaptive immune responses with some sophistication.

## INTRODUCTION

The last 20 years have seen a fundamental shift in our understanding of the roles and capabilities of neutrophils. We are now aware that they can live for much longer than previously thought (up to 5 or 6 days *in vivo* [1]); that they move into lymphatics, both during inflammation and as part of regular homeostatic control [2–8]; that they can have reparative as well as pro-inflammatory roles [9–14]; and that they influence outcomes in chronic inflammatory and autoimmune diseases, as well as infections [15–17]. As a result, we now recognise neutrophils as able to influence and shape adaptive immunity in tissues and in lymph nodes, in both suppressive and activatory manners (reviewed in [18, 19]). For example, neutrophils restrain Th2 inflammation during asthma [20], suppress dendritic cell (DC) migration to lymph nodes [21], suppress T cell proliferation in sepsis via Mac-1 [22], and limit humoral responses in lymph nodes [23]. In contrast, they can also directly present antigen to T cells via MHC class II [24], cross-present to CD8^+^ T cells [25], directly stimulate T cell proliferation in response to superantigen [26], promote type 2 responses in the lung [27], increase DC maturation and co-stimulatory molecule expression [16, 28–30] and promote T cell function during influenza infection [31, 32].

A particular relationship between neutrophils and the IL-17-producing Th17 subset of CD4^+^ T cells is now evident. The two cells chemoattract each other to the site of inflammation through production of CCL20/22, IL-17 and IL-8 [33], and the production of IL-17A and IL-17F by Th17 cells increases epithelial cell release of G-CSF and CXCL8, increasing neutrophil migration and activation [34]. Following release of extracellular traps, the surviving neutrophil cytoplast structures can indirectly induce Th17 differentiation in the lung during severe asthma [35]. Neutrophils also produce the Th17 differentiation cytokine IL-23 [36], their elastase release processes DC-derived CXCL8 into a Th17-promoting form [37], and *in vitro* they can induce Th17 differentiation directly [38]. However, the mechanisms through which this occurs largely remain unknown.

A major component of neutrophil secondary granules [39, 40] and extracellular traps [41] is the short helical antimicrobial host defence peptide (HDP) cathelicidin (human hCAP-18/LL-37, mouse mCRAMP). It can also be produced in to a lesser extent by macrophages, mucosal epithelium, eosinophils and mast cells. Release of HDP, such as cathelicidin, is a critical part of the first line innate immune response to infection [42, 43] and it is antibacterial, antiviral, antifungal and immunomodulatory [44–46], with potent ability to modulate the local innate and adaptive immune response. Amongst other effects, it can act as a chemoattractant for immune cells [47, 48], promote protective inflammatory responses and modulate cell death [49, 50], induce wound healing, re-epithelialization and re-endothelialization [51, 52], allow the take-up of self-RNA and production of type one interferons by plasmacytoid DC [53, 54] and inhibit class switching in B cells [55].

We and others have previously shown cathelicidin to directly and indirectly affect T cell function; it induces DC to prime increased proliferation and pro-inflammatory cytokine production by CD8^+^ T cells [56], is a chemoattractant for T cells [47], and, in psoriasis, it is recognised directly as an autoantigen by CD4^+^ T cells [57].

The outcome of neutrophil cathelicidin-CD4^+^ T cell interaction is largely uncharacterised. In this project we set out to investigate how cathelicidin affected T cell differentiation. We report that cathelicidin is a potent T cell differentiation factor, which induces Th17 and suppresses Th1 differentiation during inflammation, as well as selectively suppressing death of IL-17 producing cells. We show that this is partially dependent on both the aryl hydrocarbon receptor and the presence of TGF-β1 and propose this as a major mechanism by which neutrophils can direct T cell behaviour during inflammatory disease.

## RESULTS

### The host defence peptide cathelicidin induces IL-17A production by CD4^+^ T cells

In order to determine how cathelicidin specifically affects differentiation of T helper cell subsets, we exposed splenic single cell suspensions from C57BL/6J mice to Th1-driving, Th2-driving and Th17-driving conditions. Some wells were exposed to the murine cathelicidin mCRAMP at 2.5 μM, which is a physiologically relevant concentration (with cathelicidin in airway secretions from healthy newborns averaging 2 μM and from those with pulmonary infections up to 6.5 μM [58])

Analysis of cytokine production by intracellular flow cytometry revealed that addition of cathelicidin had no impact on IFN-γ producing CD4^+^ T cells under Th1-driving conditions. We also noted no difference in IL-4 producing CD4^+^ T cells in Th2-driving conditions, indicating that cathelicidin does not affect Th1 or Th2 differentiation in this system. However, we noted a consistent increase in the frequency of IL-17A producing CD4^+^ T cells following cathelicidin exposure under Th17-driving conditions (Fig1A). The total count of CD4^+^ cells in culture did not change significantly, from a mean of 33.5 +/- 4.8 x 10^3^ CD4^+^ T cells in control samples to 38.1 +/- 5.9 x 10^3^ cells in cathelicidin-exposed samples. Further, we noted that under non-polarising T cell activation conditions (αCD3/αCD28 stimulation), cathelicidin induced no increase in IL-17A production (Fig.1B), indicating this peptide is unable to act alone, instead boosting pathways which have already been triggered.

**Figure 1:**
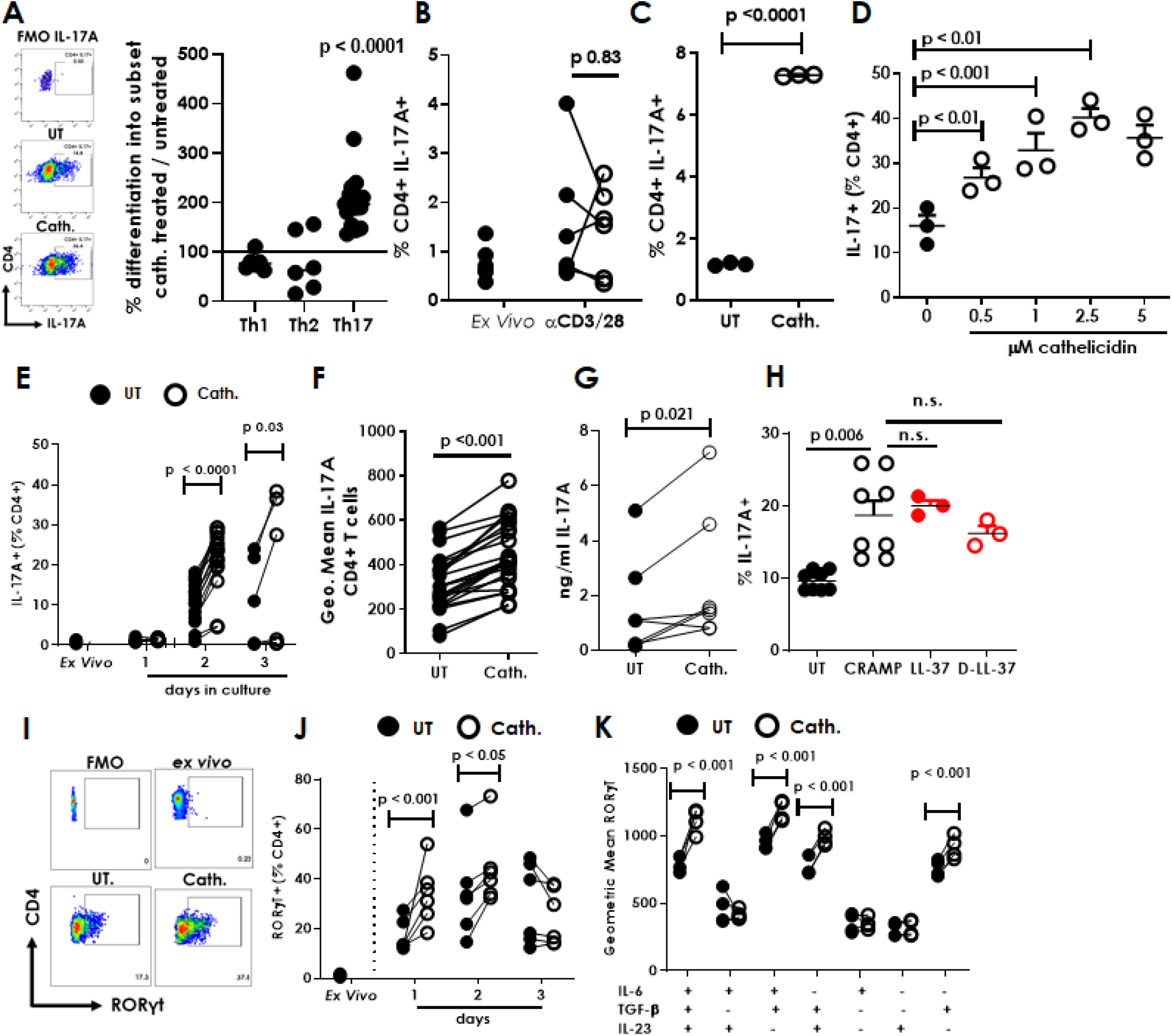
The antimicrobial peptide cathelicidin induces IL-17 production by CD4^+^ T cells. Splenocytes isolated from C57BL/6J mice were cultured in Th1-driving conditions for 4 days, Th2-driving for 4 days or Th17-driving for 2 days in the presence or absence of 2.5μM murine cathelicidin. (A,B) Expression of cytokines was determined by intracellular flow cytometry. (C) sorted CD4^+^ T cells were cultured alone in Th17-driving conditions with 2.5μM cathelicidin. (D) Production of IL-17A following increasing doses of cathelicidin (at 48 hours) and (E) production over increasing days in culture was assessed. (F) Geometric mean of IL-17A expression was also assessed on day 2 of culture. (G) After 3 days in culture the culture medium was collected and ELISA used to quantify IL-17A protein. (H) Alternative cathelicidins were used – human LL-37 and the D-enantiomer D-LL37. (I) Representative flow cytometry plots showing RORγt expression - (J) RORγt expression increased over time following cathelicidin exposure and (K) in the presence of TGF-β. Data shown are individual mice used in separate experiments, with line at median. Statistical significance was determined using a two-way repeated measures ANOVA with a Bonferroni multiple comparison post-test (E, J), a two-way ANOVA with Bonferroni correction (K), a paired t-test (B, C, F, G) or a one-way ANOVA with a Dunnett’s multiple comparison post-test (D. H).

To determine whether cathelicidin acted on CD4^+^ T cells directly, sorted cells were exposed to cathelicidin under Th17-driving conditions (Fig.1C). IL-17A production was significantly lower in sorted CD4^+^ T cell cultures compared to whole splenic cultures, but was significantly increased by cathelicidin – strikingly by a far greater amount than in whole splenic cultures, with an average 6.2-fold increase (Fig.1C).

Further investigation with whole splenic cultures revealed that this increase was dependent on both the concentration of cathelicidin present (Fig.1D) and the time in culture (Fig. 1E); cathelicidin exposure led to a 2.5 fold increase in IL-17A production by day 2 of culture, with further increases seen on day 3. Not only the frequency of IL-17A^+^ cells but also the intensity of IL-17A expression (Fig. 1F) was increased following cathelicidin exposure, with consequent increased detection of IL-17A by ELISA in cell culture supernatant on day 2 (Fig. 1G).

Cathelicidin has multiple receptors including P2XR and FPR2 [47, 59–61]. It can also enter cells including plasmacytoid DC, fibroblasts, CHO epithelial cells and bladder carcinoma cells without a receptor, via lipid-raft dependent endocytosis [54, 62]. To determine whether cathelicidin’s increase in IL-17A was receptor-dependent, we used the human cathelicidin LL-37 and the matched D-enantiomer LL-37. LL-37 induced the same increase in IL-17A in mouse cells as the murine cathelicidin mCRAMP did; in addition, the D-form peptide D-LL-37 induced a strong increase in IL-17 (UT cells 9.45 +/- 0.47 %, D-LL-37 16.2 +/- 1.01%) (Fig.1H). These results suggest that cathelicidin is functioning to modify CD4^+^ T cell phenotype in a receptor independent manner.

Th17 cells are characterised by expression of the transcription factor RORγt [63, 64]. In keeping with this, cathelicidin also induced strong expression of RORγt in CD4^+^ T cells following 24 hours in culture (Fig.1I,J).

Our Th17-driving conditions include αCD3 and αCD28 antibodies, and recombinant IL-6, TGF-β1 and IL-23. To examine whether cathelicidin enhances any of these mediators specifically, we assessed its impact on RORγt expression in the presence of each mediator individually. Cathelicidin induced increases in RORγt expression only in the presence of TGF-β1 (Fig. 1K), with no increases seen in cells treated with IL-6 alone, IL-23 alone or IL-6 and IL-23 in combination (Fig.1K). Cathelicidin enhanced RORγt in the presence of TGF-β1 alone, but peak expression following cathelicidin exposure occurred in the presence of IL-6 as well as TGF-β1. Together, these results reveal that the antimicrobial peptide cathelicidin significantly promotes Th17 differentiation and IL-17A production in a receptor-independent and TGF-β1 dependent fashion.

### Cathelicidin induces IL-17F-producing but not IL-22-producing cells

Th17 cells produce a number of pro-inflammatory cytokines including IL-17A, IL-17F and IL-22 [65–67]. We wondered whether cathelicidin would increase expression of these other cytokines in addition to IL-17A. Analysis of cathelicidin-exposed CD4^+^ T cells showed, surprisingly, that cathelicidin did not boost IL-17A single positive (SP) cells (Fig.2A, B). Instead, IL-17F single positive and A+F+ double positive cells were significantly increased by treatment (Fig.2 C, D). Interestingly, production of IL-22 was not altered significantly by cathelicidin (Fig.2E).

**Figure 2:**
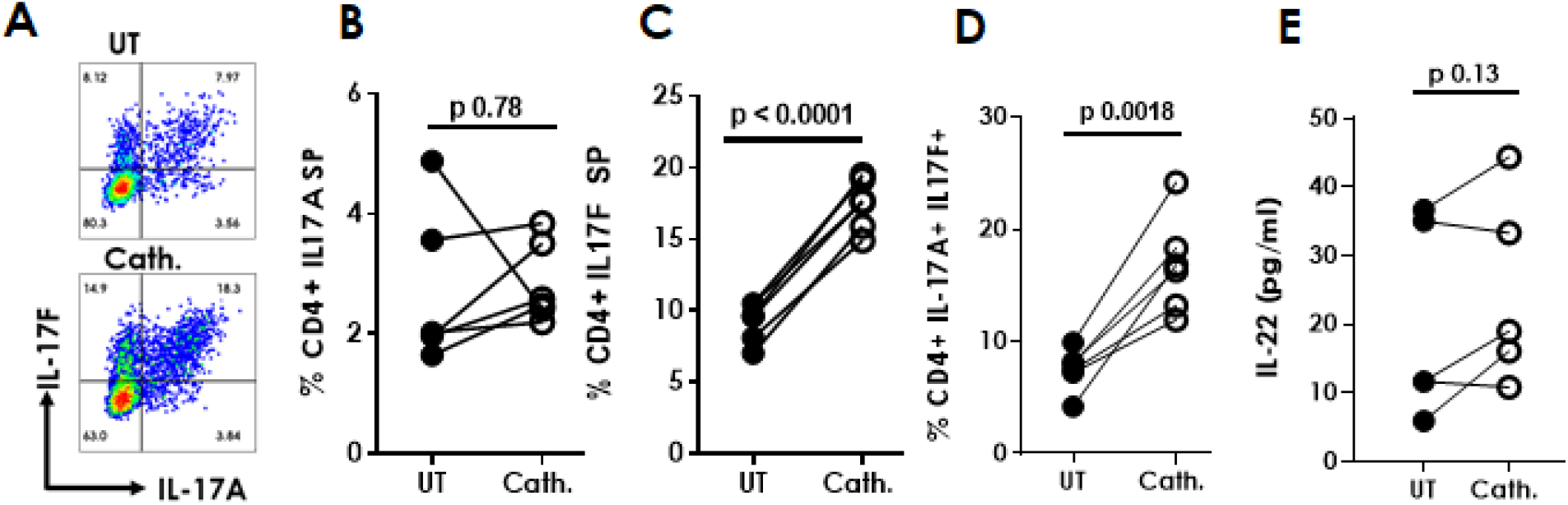
Cathelicidin induces IL-17F-producing but not IL-22-producing cells. Splenocytes isolated from C57BL/6J mice were cultured in Th17 conditions for 2 days in the presence or absence of 2.5μM murine cathelicidin. (A-D) Expression of IL-17A and F was determined by intracellular flow cytometry. (E) IL-22 production was assessed by ELISA after two days in culture. Data shown are individual mice used in separate experiments. B-E are analysed by paired two-tailed T tests. SP = single positive.

### Cathelicidin-mediated enhancement of IL-17A production, but not IL-17F, occurs via the aryl hydrocarbon receptor

Next, to understand more about the induction of Th17 cells by cathelicidin, we performed gene expression analysis on sorted CD4^+^ T cells exposed to Th17-driving conditions in the presence or absence of 2.5 μM cathelicidin for 24 hours, using a Nanostring mouse immunology chip. Following cathelicidin exposure, we noted a consistent, significant decrease in the expression of a number of genes which suppress the induction or conversion of cells into the Th17 subset [68–71] including *Socs3, Stat1, Irf8, Bcl3* and *Ikzf4* (Fig. 3A, B).

**Figure 3:**
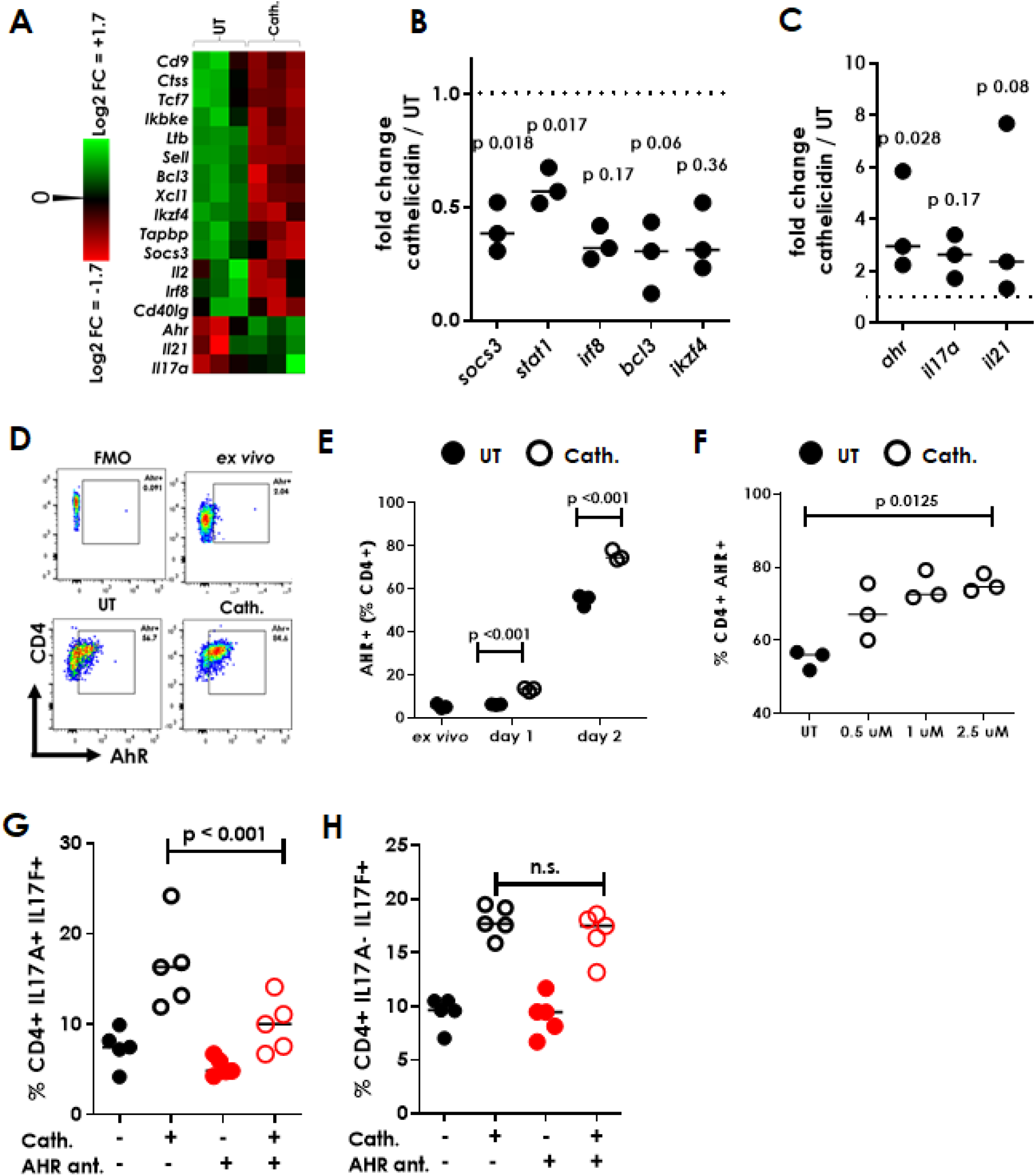
Cathelicidn induces Th17 cells via AhR up-regulation. CD4^+^ T cells isolated from C57BL/6J mice were cultured in Th17-driving conditions for 24 hours in the presence or absence of 2,5μM murine cathelicidin. (A-C) gene expression differences were assessed by a Nanostring mouse immunology chip. (D) representative flow cytometry plots showing aryl hydrocarbon receptor staining, which was assessed (E) overtime and (F) with increasing doses of cathelicidin. (G, H) IL-17A and F production was assessed following blockade of AhR with the antagonist CH223191 (ant.). Data shown are individual mice used in separate experiments, with line at median. Statistical analyses: (B, C) - paired two-tailed t tests with Bonferroni multiple comparison post-test, E - two-way repeated measures ANOVA with a Bonferroni multiple comparison post-test; F - one-way ANOVA with a Dunnett’s multiple comparison post-test; G, H two-way ANOVA with Tukey’s multiple comparison post-test.

In keeping with our initial observations, cathelicidin exposure increased expression of the genes coding for IL-17A and the Th17 cytokine IL-21 (Fig.3A, C). Intriguingly, cathelicidin also led to a large increase in expression of the aryl hydrocarbon receptor (AHR) gene *Ahr* (Fig.3A, C). Flow cytometric analyses subsequently confirmed an increase in AHR protein following cathelicidin exposure under Th17-driving conditions (Fig.3 D, E), which was dependent on time of exposure to cathelicidin and on concentration of the peptide present (Fig.3 E, F). The increase in AHR expression by cathelicidin was significant after one day in culture (Fig.3E), earlier than the IL-17 increase was noted.

AHR is a known Th17 differentiation factor [66, 68, 72] and so we hypothesised that the cathelicidin-mediated increase in IL-17 was a result of its enhanced expression. To test this, we used the specific AHR antagonist CH223191 [73]. Use of CH223191 abolished the increase in IL-17A^+^F^+^ double-producing cells induced by cathelicidin, indicating that cathelicidin’s induction of these Th17 cells is dependent on AHR (Fig.3G). Interestingly, single IL-17F-producing cells were increased by cathelicidin, as previously shown (Fig.2C), but this increase was not significantly altered by AHR antagonism (Fig.3H). These results indicate that there are both AHR-dependent and independent mechanisms through which cathelicidin promotes Th17 differentiation, with AHR being critical for the enhancing of all IL-17A producing cells, but not the increase in cells producing only IL-17F.

### Cathelicidin suppresses Th1 differentiation in the presence of TGF-β1

Gene expression analysis also revealed a consistent decrease in Th1-associated genes, including *Il2, Irf8, Stat1*, and *Xcl1* (Fig.3A) following 24 hours of cathelicidin exposure in cells differentiating under Th-17 driving conditions. Expression of IL-2 protein was confirmed to be reduced following cathelicidin exposure on day 2 of culture (Fig.4A), with a striking total suppression of IL-2 production by CD4^+^ T cells, as measured by ELISA. Expression of the Th1-associated transcription factor Tbet (Fig.4B) and production of the key Th1 cytokine IFN-γ (Fig.4C, D) were also both strongly suppressed by cathelicidin. These results are in contrast to figure 1A, in which cathelicidin had no impact on IFN-γ production in Th1 driving conditions. Th17-driving conditions do suppress IL-2 production and Th1 commitment [74, 75] and we proposed that cathelicidin boosts this suppression further. In keeping with this, we determined that cathelicidin had no impact on Tbet expression in activating non-lineage driving conditions (αCD3, αCD28) but only when TGF-β1 was present (Fig.4E) – and the presence of TGF-β1 alone was sufficient for cathelicidin to induce this suppression. This demonstrates a further enhancement of TGF-β1’s action by cathelicidin.

**Figure 4:**
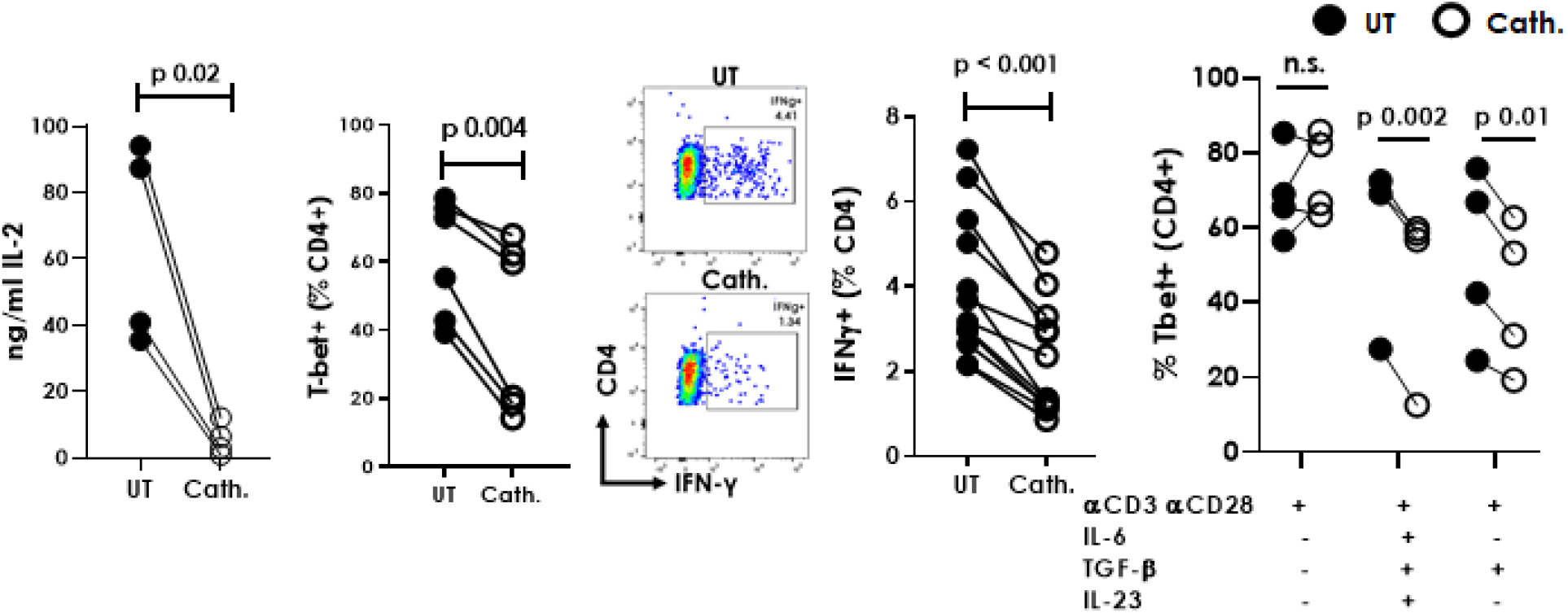
Cathelicidin suppresses Th1 differentiation in the presence of TGF-β1. Splenocytes isolated from C57BL/6J mice were cultured in Th17-driving conditions for 48 hours in the presence or absence of 2.5μM murine cathelicidin. (A) production of IL-2 was assessed by ELISA on day 2 of culture. (B-D) Expression of Tbet and IFN-γ were assessed by flow cytometry. (E) Tbet expression was assessed following incubation with individual cytokines in the presence of cathelicidin. Data shown are individual mice used in separate experiments and analysed by paired two-tailed t tests.

### Cathelicidin promotes survival of Th17 cells but not Th1 cells

We have shown that cathelicidin promotes increased differentiation of Th17 cells with suppression of Th1 cells in the presence of TGF-β1. We next investigated whether, in addition to modulated differentiation of CD4^+^ T cells, cathelicidin also led to differential survival or proliferation of particular cell subsets. Cathelicidin has been previously shown to induce death of certain T cell subsets, including T regulatory cells and CD8^+^ T cells, although CD4^+^ T effector cells were not affected [76, 77]. Our Nanostring gene expression analysis revealed a significant down-regulation of *Fasl* and consistent but not significant down-regulation of *Fas* in sorted CD4^+^ T cells 24 hours following cathelicidin exposure (Fig.5A). Subsequently, to determine whether this led to altered rates of death of CD4^+^ T cells, we measured annexin V staining and uptake of propidium iodide, which together distinguish between live, apoptotic and necrotic cells (Fig.5B). Using this method, we noted that cathelicidin significantly suppressed death of CD4^+^ T cells, with effects seen by the first day of culture (Fig.5 B,C).

**Figure 5:**
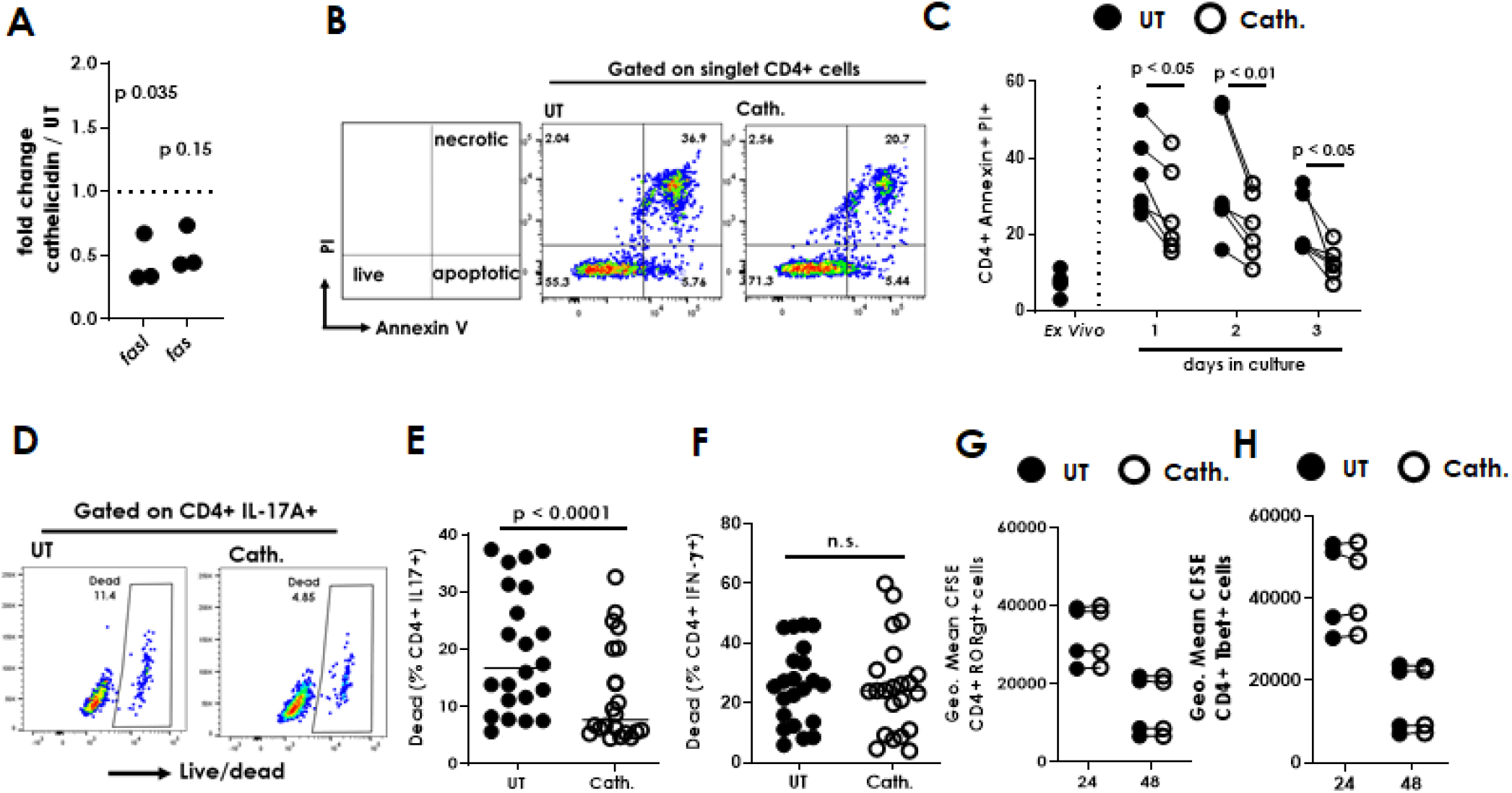
Cathelicidin protects Th17 cells but not Th1 cells from death. CD4^+^ T cells isolated from C57BL/6J mice were cultured in Th17-driving conditions for 24 hours in the presence or absence of 2.5μM murine cathelicidin. (A) gene expression differences were assessed by Nanostring mouse immunology chip. Cell death was assessed in culture by (B-C) annexin / PI staining and flow cytometry and (D-F) by binding of a fixable live-dead marker. (G-H) T cells were labelled with CFSE and proliferation assessed following two days in culture. Data shown are individual mice used in separate experiments, with line at median. A, E and F were analysed with paired two-tailed t tests. C was analysed by a two-way repeated measures ANOVA with a Bonferroni multiple comparison post-test.

To investigate suppression of death by cytokine producing cells in particular, uptake of fixable live dead dyes was assessed (Fig.5D). Interestingly, cathelicidin exposure suppressed death of IL-17A-producing CD4^+^ T cells (Fig.5D, E) but not IFN-γ-producing cells (Fig.5F), in the same samples, under Th17-driving conditions. To our knowledge, this is the first demonstration of a neutrophil peptide, or of any antimicrobial peptide, increasing survival of T cells. These data raise the possibility that neutrophils may have sophisticated impacts on developing and skewing T cell immunity during inflammatory disease.

It was also possible that cathelicidin induces increased proliferation of Th17 cells but not Th1 cells, and that this contributes to their increased frequency in culture. CD4^+^ T cells labelled with CFSE were observed following exposure to cathelicidin. Although CFSE intensity halved from day 1 to 2 of culture, as the cells divided (Fig.5G,H), there was no difference in the proliferation of Th17 or Th1 cells following exposure to cathelicidin, indicating increased division is not a mechanism by which cathelicidin increases frequency of Th17 cells.

### Mice lacking cathelicidin cannot increase IL-17 production in response to inflammation

Next, we questioned whether our results extended *in vivo*, and whether T cells develop normally in cathelicidin knockout (*Camp^tm1Rig^*, KO) mice. Firstly we assessed whether KO mice had altered Th17 cell numbers in the steady state. Analysis of *ex vivo* T cell cytokine production in multiple organs revealed no differences between KO and wildtype C57BL/6JOlaHsd (WT) mice (Fig.6A,B) – indicating that cathelicidin is not required for development of Th17 cells in the steady state.

**Figure 6:**
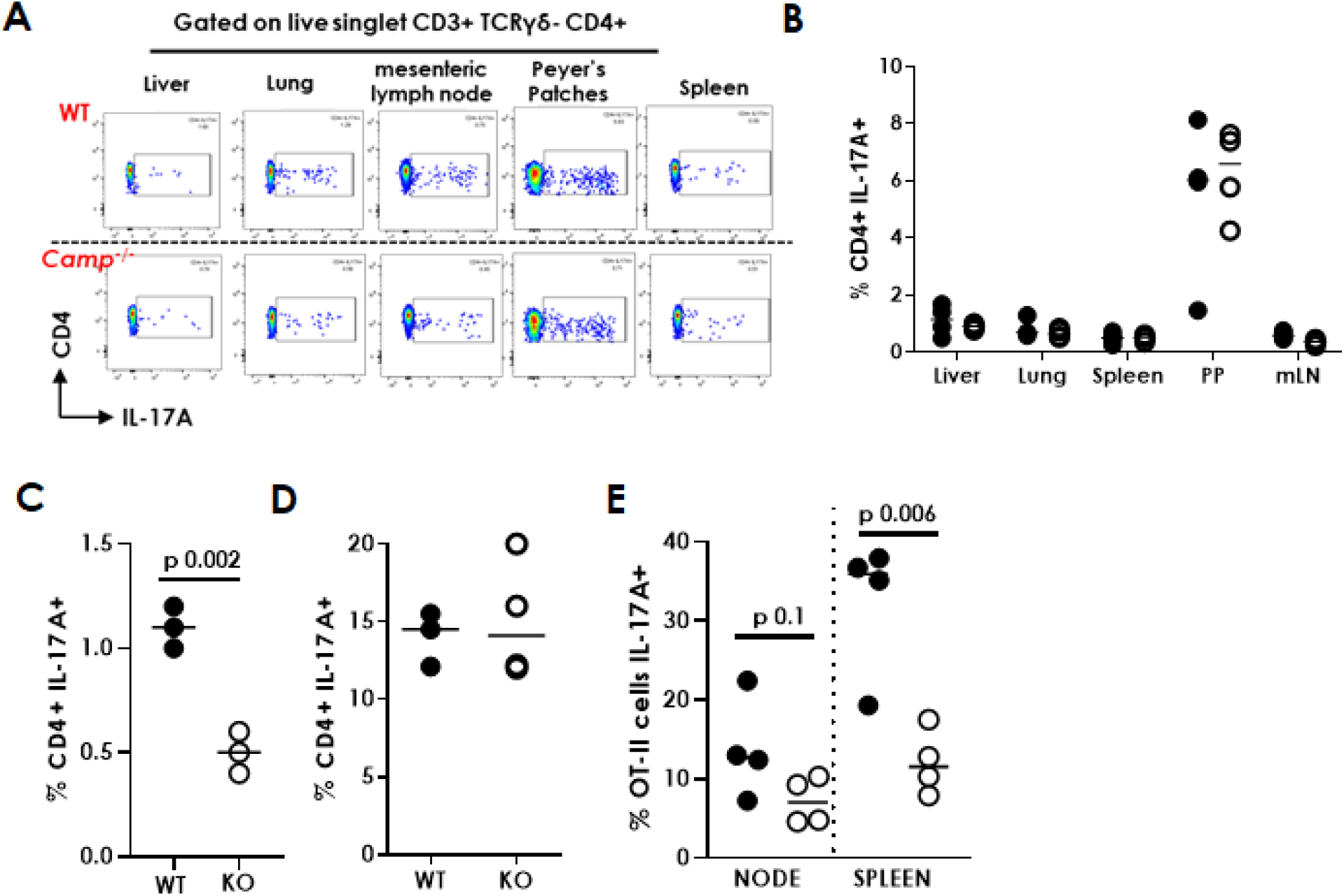
Mice lacking cathelicidin cannot increase IL-17 production in response to inflammation. Wildtype (WT) C57BL/6/JOIaHsD mice and fully backcrossed cathelicidin knockout (*Camp^-/-^*, KO) mice were culled and (A, B) organs removed and IL-17 production assessed *ex vivo* by flow cytometry. (C) mice were inoculated with 12.5μg heat-killed *Salmonella typhimurium* and draining lymph nodes removed seven days later-production of IL-17 was measured by flow cytometry. (D) Splenic T cells from WT or KO mice were stimulated in Th17-driving conditions for 2 days and IL-17 production measured by flow cytometry. (E) WT and KO cells were injected intravenously with 5M OT-II cells and 24 hours later sub-cutaneously with ovalbumin in complete Freund’s adjuvant. On day 7 spleens and lymph nodes were removed and IL-17 production by donor cells assessed by flow cytometry. Data shown are individual mice with line at median. C and E were analysed by two-tailed t tests.

We hypothesised that while cathelicidin has no impact on normal T cell development, it may affect Th17 differentiation during inflammation, when it is released by neutrophils and other cells. To investigate the impact of cathelicidin in the context of inflammation, we used a vaccination model. WT and KO mice were inoculated with heat killed *Salmonella typhimurium* (HKST) into the top of each hind paw. Seven days later draining popliteal lymph nodes were removed and IL-17A quantified by flow cytometry. KO lymph node CD4^+^ T cells were unable to produce IL-17 to the same level as WT T cells (Fig.6C), confirming that cathelicidin promotes Th17 differentiation *in vivo* as well as *in vitro*.

This experiment did not tell us if the T cells themselves were inherently different in KO mice or unable to produce IL-17. To test this, we firstly stimulated KO splenic T cells *in vitro*. In the presence of Th17-driving cytokines, CD4^+^ T cells from KO mice could produce IL-17 normally (Fig.6D). Secondly, we performed an adoptive transfer of OT-II CD4^+^ T cells into WT and KO mice and immunised the mice with ovalbumin in complete Freund’s adjuvant. Analysis of donor T cells in draining inguinal lymph nodes and spleens 7 days later showed significantly lower IL-17A production by donor T cells introduced into KO mice (Fig.6E). These two experiments combined led us to conclude that a) cathelicidin is not required for normal development of Th17 cells or for baseline production of IL-17 but b) in the context of inflammation, cathelicidin enhances Th17 differentiation and IL-17 production. They also demonstrated that our *in vitro* observation that cathelicidin is a Th17 differentiation factor was relevant *in vivo*.

### Cathelicidin is released in the lymph nodes by neutrophils

Repeating our *in vitro* culture experiments with Th17-driving medium, we found that cathelicidin needed to be present in the first 24 hours of culture in order to induce Th17 differentiation (Fig.7A), with its addition to activated cells leading to no differences in IL-17 production. This suggests that, *in vivo*, it would only promote Th17 differentiation if present in lymph nodes during the antigen presentation process, and would not have an enhancing effect in the tissue if interacting with T cells which are already activated and differentiated.

**Figure 7:**
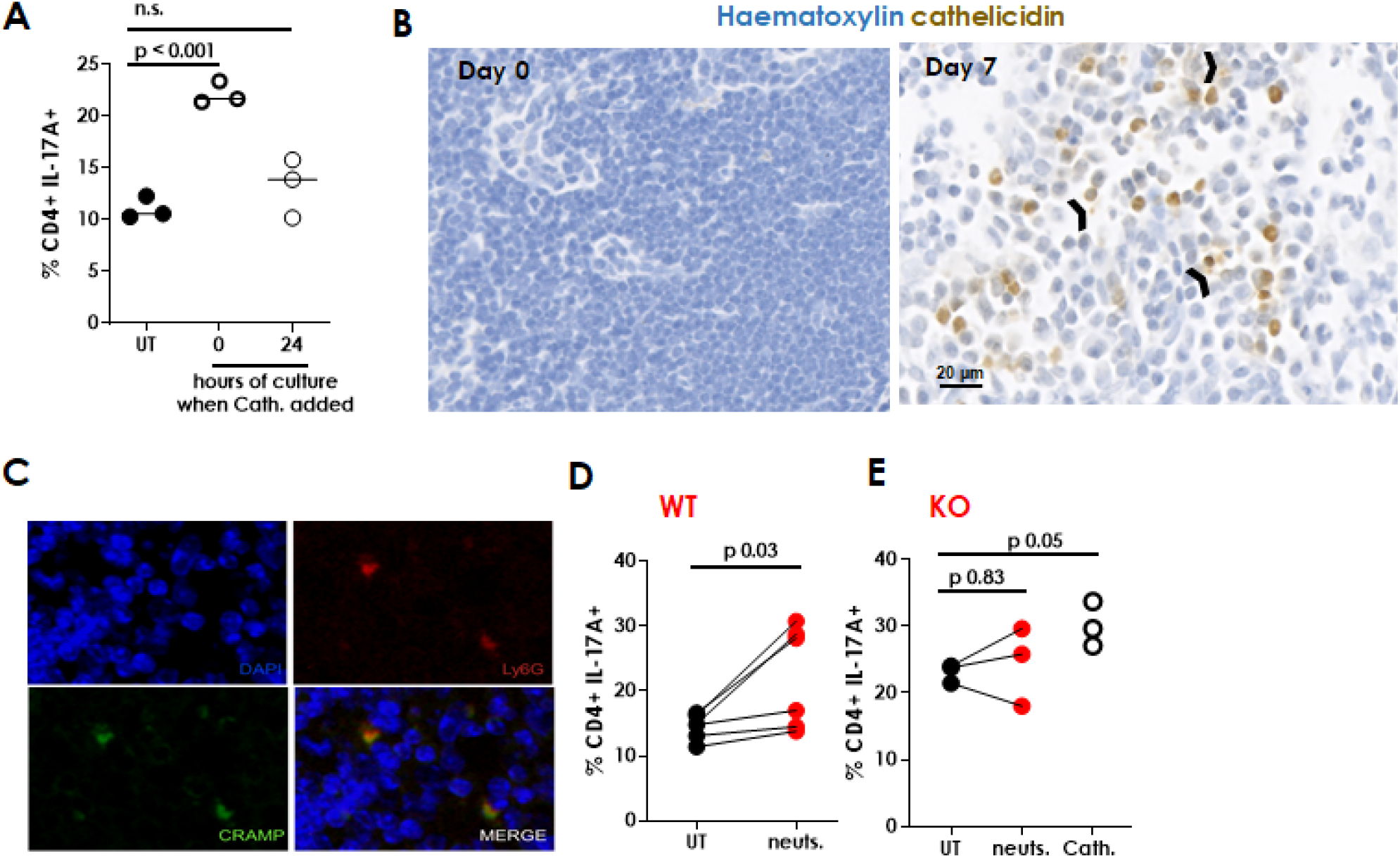
Cathelicidin is released in the lymph nodes by neutrophils. (A) IL-17 production was assessed by flow cytometry following cathelicidin being added to murine Th17 cultures on day 0 or day 1. (B, C) Wildtype C57L/6JOIaHsD mice were inoculated with 12.5 μg heat-killed *Salmonella typhimurium* (HKST) and draining lymph nodes removed up to 7 days later. Lymph nodes on day 0 and 7 following HKST inoculation were assessed for cathelicidin (B, arrow heads indicate cathelicidin not associated with cells) and neutrophil numbers (C). Neutrophils were isolated from bone marrow of (D) wildtype (WT) and (E) fully backcrossed cathelicidin knockout (KO) mice and added to Th17-generating cultures of splenic CD4^+^ T cells for 48 hours before T cell cytokine production was assessed by flow cytometry. Data shown are individual mice with line at median (A), representative images from three separate experiments (B,C), or individual mice (D,E). Cath= 2.5 μM cathelicidin. A was analysed by a one-way ANOVA with a Dunnett’s multiple comparison post-test. D and E were analysed by paired two-tailed t tests.

We did not know whether or when cathelicidin was produced in the lymph nodes. To determine this, we examined lymph nodes over a time course of HKST injection. Cathelicidin was not present in the steady state lymph node but appeared on day one following inoculation and was strongly present at day 7 (Fig.7B). Interestingly, we noted the presence of extracellular cathelicidin (arrow heads, Fig.7B) which had been released from cells.

Co-staining lymph node sections revealed that the vast majority of cathelicidin-expressing cells were neutrophils, with 85% of cathelicidin signal on day 7 being associated with Ly6G^+^ cells (Fig. 7C). We therefore hypothesised that neutrophils were releasing cathelicidin in the lymph nodes and enhancing Th17 differentiation. We examined whether neutrophil release of cathelicidin could induce Th17 differentiation *in vitro*. Neutrophil-induced production of IL-17 has previously been demonstrated [78]. Here, we used mouse bone-marrow isolated neutrophils from WT or KO mice, which had been activated by 30 minutes’ incubation with fMLF and cytochalasin B, to induce de-granulation. The neutrophils were included in culture wells with WT CD4^+^ T cells, at a 1:1 T cell: neutrophil ratio and in Th17-driving conditions. WT neutrophils induced IL-17 production by CD4^+^ T cells (Fig.7D) but KO neutrophils did not, although the positive control cathelicidin demonstrated that the experiment had worked as normal (Fig.7E). This data demonstrates that the increase in IL-17A production induced by de-granulating neutrophils is owing to their release of cathelicidin.

### Cathelicidin induces IL-17A production from human CD4^+^ T cells

Finally, to assess whether our observations held true for human cells, we isolated CD4^+^ T cells from peripheral blood of healthy human donors and incubated them with 2.5 μM human cathelicidin (LL-37) under Th17-driving conditions. Production of IL-17A was increased following cathelicidin exposure, measured by flow cytometry on day 8 of culture (Fig.8A) and ELISAs on days 3 and 8 (Fig.8B).

**Figure 8:**
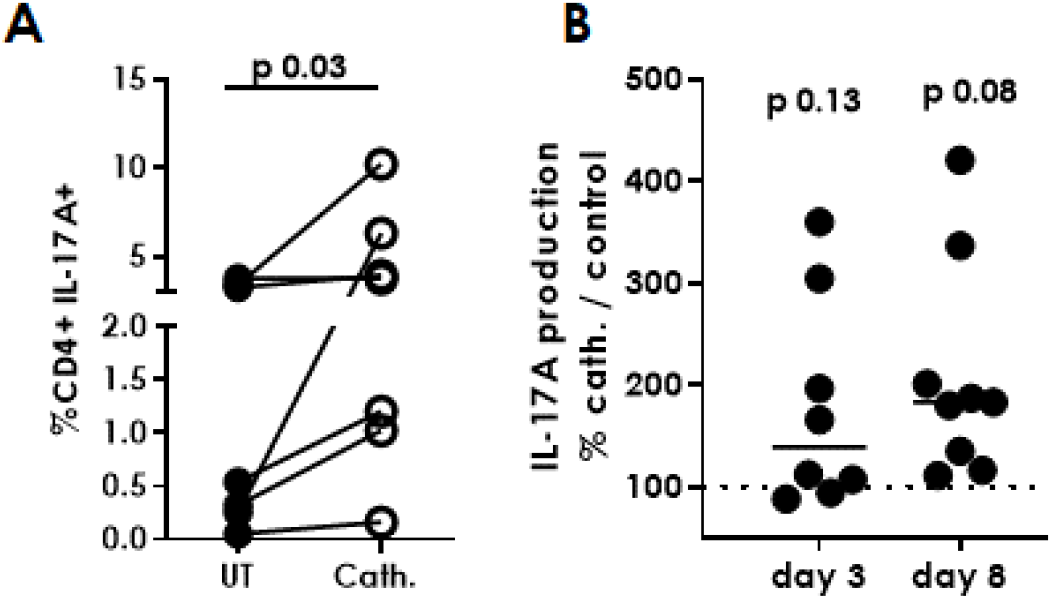
Human CD4^+^ T cells enhance IL-17 production following cathelicidin exposure. CD4^+^ T cells were isolated from peripheral blood from healthy human donors and cultured for 8 days in Th17-driving conditions, in the presence or absence of 25μM cathelicidin. IL-17A production was assessed by (A) intracellular flow cytometry on day 8 and (B) ELISA on cell culture supernatant. Data shown are individual donors with line at median. A and B are analysed by paired t tests. B on raw data before normalisation.

## DISCUSSION

We demonstrate that the mouse and human antimicrobial host defence peptide cathelicidin (LL-37/ mCRAMP) induces differentiation of Th17 cells *in vitro* and *in vivo*. Specifically, CD4^+^ T cells exposed to cathelicidin a) up-regulated AHR, RORγt, IL-17A and IL-17F b) down-regulated IL-2, IFN-γ and Tbet and c) survived in greater numbers than their untreated controls. This work extends previous studies showing that neutrophils and Th17 cells amplify each others’ responses (reviewed in [18]), providing a novel explanation for how neutrophils specifically promote one subset of T cells above others.

Cathelicidin exposure led to a 6-fold increase in production of IL-17A in sorted CD4^+^ T cells (Fig.1D), and a 2.5-fold increase in production by CD4^+^ T cells in whole splenic cultures (Fig.1E). This compares to a 2.5-fold increase in IL-17A production by the aryl hydrocarbon receptor FICZ [68] and a 3-4-fold increase in IL-17A mRNA by the same reagent [66]. These results therefore identify cathelicidin as of equal potency to one of the most important Th17 inducers known.

We show that cathelicidin specifically increases frequency of and cytokine production by those cells which produce IL-17F as well as IL-17A, with seemingly separate pathways promoted in each case. Antagonism of the AHR pathway led only to a suppression in cathelicidin’s induction of cells producing IL-17A and not in the induction of IL-17F sole producers (Fig.3G,H). Previously, triggering of AHR with FICZ led to an increase in IL-17A sole producers and IL-17A^+^F^+^ dual producers, but led to a decrease in IL-17F single positive cells [66]. This, along with our data, implies that there may be two pathways promoted by cathelicidin: one of which is AHR-dependent, and which enhances IL17A+F+ production; and one which is AHR-independent and which enhances IL-17F production alone. This may also explain the interesting lack of difference in IL-22 production by cathelicidin-treated cells, as this has also been shown to be driven by AHR ligation [66]. Single-cell transcriptome and signalling pathway analysis of whole splenic cultures treated with cathelicidin will be required to test this hypothesis, and to extend our understanding of how cathelicidin is enhancing TGF-β1 and AHR signalling.

The up-regulation of RORγt and down-regulation of Tbet induced by cathelicidin was dependent on the presence of TGF-β1 in the differentiation cocktail. TGF-β1 has been used to differentiate mouse and human Th17 cells in many studies [79] [80], and cathelicidin could not induce Th17 cells in its absence, but rather enhanced its effects. The necessity for TGF-β1 in cathelicidin’s induction of Th17 cells extends previous observations of interplay between these two mediators in other cell types [81–83]. Neutrophils can produce TGF-β in lymph nodes and in tissues. Interestingly, lymph node-neutrophil inhibition of B cell activation following inoculation is also TGF-β dependent [23]. Deposition of granule proteins in the lymph node by de-granulating neutrophils has been demonstrated previously [21] and the presence of extracellular cathelicidin in our lymph nodes following HKST inoculum supports our hypothesis that deposition of this peptide is also occurring during inflammation, alongside production of TGF-β from multiple cell types. This allows the enhanced differentiation of Th17 cells during inflammation.

Exposure of CD4^+^ T cells to cathelicidin under Th17-driving conditions led to an almost complete suppression of IL-2 production, an enhancement of that noted previously as induced by AHR signalling [74]. This observation is novel and intriguing. It would predict that T cells exposed to cathelicidin, in the presence of de-granulating or NETosing neutrophils, will proliferate less, as the available IL-2 is reduced. We do not see a difference in proliferation in our cultures after two days in culture, ruling this out as a cause for the increased number of Th17 cells present in culture. However, it will be interesting to examine longer culture times. Further, neutrophil-induced suppression of T cell proliferation is well documented [22, 84]. We suggest that the neutrophils’ release of cathelicidin, and subsequent suppression of IL-2 production, is one mechanism behind this phenomenon.

The increased survival specifically of Th17 cells following cathelicidin exposure is interesting. Neutrophil-boosted survival of any T cell subset has never previously been described. Neutrophils can release BAFF and APRIL in the lymph nodes and thus protect B cells against apoptosis [85–87], and can also enhance NK cell survival [88], meaning our novel demonstration of T cell survival fits within the context of the wider lymphocyte population.

Th17 cells are associated with a number of inflammatory and autoimmune conditions, including psoriasis, rheumatoid arthritis, and type 1 diabetes (reviewed in [89]). Our work supports previous data showing that cathelicidin is recognised by some CD4^+^ T cells as an auto-antigen during psoriasis, and those T cells which recognise cathelicidin produce more IL-17 [57]. We hypothesise then that the generation of long-lived T cells producing more IL-17 would be deleterious, and therefore that neutrophils present during these conditions could be responsible for increased disease onset and severity. In this regard, however, the specific increase in IL-17F producing cells by cathelicidin is interesting, as this cytokine has been correlated with non-pathogenicity of cells [90, 91], and the spinal-cord infiltrating Th17 cells in EAE are mostly solely producing IL-17A [91]. Future work understanding the role of neutrophil-released cathelicidin in altering the pathogenicity of Th17 cells, and in the specific induction of IL-17F, will be of great interest.

Overall, our study identifies the antimicrobial peptide cathelicidin as a key Th17 differentiation factor and defines one mechanism by which neutrophils can alter developing T cell responses in a sophisticated and specific manner.

## METHODS

### EXPERIMENTAL DESIGN

All experiments were performed three times on individual mice or donor cell cultures and then power calculations performed to determine necessary group sizes. Both male and female mice and human donors were used in experiments and no differences were noted in responses between the sexes.

Mice were housed together for *in vivo* experiments to avoid cage-specific effects.

### MICE

Wild type C57Bl/6JOlaHsd and mCRAMP knockout (*Camp^tm1Rig^*, KO) mice were bred and housed in individually ventilated cages, under specific pathogen-free conditions. Male and female mice between 6-12 weeks of age were used. KO mice were backcrossed onto wildtypes for 10 generations before use.

All animal experiments were performed by fully trained personnel in accordance with Home Office UK project licences PAF438439 and 70/8884, under the Animal (Scientific Procedures) Act 1986.

### HEALTHY HUMAN DONORS

Peripheral venous blood was collected from healthy adult volunteers under ethical agreement code AMREC 15-HV-013. Up to 160ml of blood was collected at one time into sodium citrate and was then processed immediately. Ficoll Paque Plus (GE Healthcare #GE17-1440-02) was used to isolate mononuclear cells by mixing with freshly isolated blood (1:1 diluted in PBS) and spinning in LeucoSep tubes (Grenier #227289) for 15 minutes at 1000g, with brake at 0. CD4^+^ T cells were then isolated from the cell layer using the EasySep™ human CD4^+^ T cell isolation kit (StemCell Technologies #17952), according to manufacturers’ instructions.

### HUMAN TH17 CULTURES

Human CD4^+^ T cells were plated at 1×10^5^ cells/well in a round-bottom 96-well plate in complete medium (RPMI, 10% fetal calf serum, 10 units/mL penicillin, 10 μg/mL streptomycin and 2 mM L-glutamine, all supplied by Gibco, ThermoFisher UK). Cytokines added were: 10ng/ml TGF-β1 (Biolegend #580702), 100ng/ml IL-6 (Biolegend #570802), 30ng/ml IL-1β (Biolegend #479402), 30ng/ml IL-23 (Biolegend #574102) and 2.5μl/ml anti-CD3/23/2 ImmunoCult T cell activator (StemCell Technologies #10970). Medium was changed on day 3 and samples were assessed by flow cytometry and ELISA on day 8.

### PEPTIDES

Synthetic mCRAMP (GLLRKGGEKIGEKLKKIGQKIKNFFQKLVPQPEQ) and LL-37 (LLGDFFRKSKEKIGKEFKRIVQRIKDFLRNLVPRTES) were custom synthesized by Almac (Penicuik, Scotland) using Fmoc solid phase synthesis and reverse phase HPLC purification. Peptide identity was confirmed by electrospray mass spectrometry. Purity (>95% area) was determined by RP-HPLC and net peptide content determined by amino acid analysis. D-enantiomer LL-37 was a kind gift from Professor Peter Barlow, Edinburgh Napier University. Lyophilized peptides were reconstituted in endotoxin free water at 5 mg/ml. Reconstituted peptides were tested for endotoxin contamination using a Limulus Amebocyte Lysate Chromogenic Endotoxin Quantitation Kit (Thermo Scientific, UK #88282).

### MURINE TISSUE AND SINGLE CELL PREPARATIONS

Mice were culled by rising concentrations of CO_2_, followed by cervical dislocation. Single cell preparations of spleens, lymph nodes and Peyer’s patches were achieved by mashing the tissues through a 100 μM strainer and washing with PBS. One lung lobe was minced and enzymatically digested with 1mg/ml collagenase VIII (Sigma Aldrich, #C2139) for 20 mins at 37°C, with shaking before the cell suspension was passed through a 100 μM filter. To isolate liver leukocytes, one lobe was passed through a 100 μM strainer and leukocytes isolated by Percoll separation, as previously described [92]. In all cases red blood cells were lysed using RBC Lysis Buffer, as per the manufacturer’s instructions (BD Biosciences, #555899).

### ISOLATION OF BONE MARROW-DERIVED NEUTROPHILS

Femurs were removed and marrow flushed out with complete medium (RPMI, 10% fetal calf serum, 10 units/mL penicillin, 10 μg/mL streptomycin and 2 mM L-glutamine, all Gibco, ThermoFisher UK). Single cell suspensions were prepared by passing the cells through a 19G needle. Neutrophils were isolated using the EasySep™ Mouse Neutrophil Enrichment Kit (StemCell Technologies, #19762), as per the manufacturer’s guidelines. Neutrophils were activated by 30 minutes’ incubation with 100 nM fMLF (N-formylmethionine-leucyl-phenylalanine, Sigma Aldrich #F3506) and 10 μM cytochalasin B (Sigma Aldrich #C2743) before being well washed twice.

### *IN VITRO* T HELPER CELL SUBSET DIFFERENTIATION

Whole splenocytes were prepared as previously described. CD3^+^ T cells were isolated from whole splenocytes using the EasySep™ Mouse T Cell Isolation Kit (StemCell Technologies, #19851), as per the manufacturer’s guidelines. CD4^+^ and CD8^+^ T cells were isolated from whole splenocytes by FACS sorting on a BD Biosciences Aria sorter using a 70μM nozzle.

200,000 cells were plated per well of round-bottom 96-well plates in complete medium (RPMI, 10% fetal calf serum, 10 units/mL penicillin, 10 μg/mL streptomycin and 2 mM L-glutamine, all supplied by Gibco, ThermoFisher UK), with the correct combination of cytokines and neutralizing antibodies as follows. Th1 cells were differentiated for 4 days in the presence of plate-bound αCD3 (5 μg/ml; Biolegend, #100339), rIL-12 (25 ng/ml; Biolegend, #575402), rIL-18 (25 ng/ml; Gibco, #PMC0184) and rIL-2 (10 U/ml; Biolegend, #575402), with or without 2.5 μM mCRAMP. Th2 cells were cultured for 4 days with plate-bound αCD3 (5 μg/ml; Biolegend, #100339), rIL-4 (4 ng/ml; Peprotech, #214-14), rIL-2 (40 U/ml; Biolegend, #575402), αIL-12 (5 μg/ml; Biolegend, #505307) and αIFNγ (5 ug/ml; Biolegend, #505812), with or without 2.5 μM mCRAMP. Th17 cells were differentiated in the presence of plate-bound αCD3 (5 μg/ml; Biolegend, #100339), rIL-6 (20 ng/ml; Biolegend, #575706), rIL-23 (20 ng/ml; Biolegend, #589006) and rTGFβ (3 ng/ml; Biolegend, #580706), with or without 2.5 μM mCRAMP for 1 to 3 days. αCD28 (1 μg/ml; Biolegend, #102115) was added to cultures of pure CD4^+^ T cells. In some experiments neutrophils, isolated as above, were incubated in CD4^+^ T cell cultures at a 1:1 neutrophil: T cell ratio.

### FLOW CYTOMETRY

Cells were stained for surface markers for 30 mins at 4°C, protected from light. Intracellular cytokines were assessed by incubating cells for 4 hours at 37°C with Cell Stimulation Cocktail containing protein transport inhibitors (eBioscience, #00-4970-03). Cells were fixed, permeabilised and stained for cytokines using BD Cytofix/Cytoperm (BD Biosciences, #554722), as per the manufacturer’s guidelines. Cells were fixed, permeabilised and stained for transcription factors using the True-Nuclear Transcription Factor Buffer Set, as per the manufacturer’s instructions (Biolegend, #424401). Cell viability was assessed by flow cytometry using the Annexin-V-FLUOS Staining Kit, as per the manufacturer’s guidelines (Roche, #11 858 777 001) or fixable live/dead yellow (ThermoFisher #L34959). To assess proliferation, cells were stained with carboxyfluorescein succinimidyl ester at 1:2000 dilution for 10 minutes (Invitrogen, #C34554) before culture. Proliferation analysis by dye dilution was performed by flow cytometry. Samples were analysed using a Fortessa LSR II (BD Biosciences) and FlowJo software (Treestar).

### ANTIBODIES (MOUSE)

CD4 (clone GK1.5, Biolegend, #100453); CD8 (53-6.7, BD Biosciences, #563786); IFNγ (XMG1.2, Biolegend, 505825); IL-17A (TC11-18H10.1, Biolegend, #506938); IL-22 (POLY5164, Biolegend, #516411); RORγT (B2D, eBiosciences, #12-6981-80); TBET (4B10, Biolegend, #644805); AHR (4MEJJ, eBiosciences, #25-5925-80)

### ANTIBODIES (HUMAN)

CD4 (clone OKT4, eBioscience #25-0048-42), IL-17A (eBio64DEC17, eBioscience #12-7179-42).

### NANOSTRING

Mouse Th17 cultures were set up as previously described. DAPI^-^γδ^-^CD4^+^ T cells were sorted using a BD FACSAria™ II (BD Biosciences) on day 1. RNA was extracted immediately using the Qiagen RNeasy Mini Kit (Qiagen, #74104), as per the manufacturer’s guidelines. Multiplex gene expression analysis (Mouse Immunology Panel) was performed by HTPU Microarray Services, University of Edinburgh. Data analysis was performed using nSolver 4.0 and nCounter Advanced Analysis software. Genes with a log2 fold change of 1 or more and a p value of 0.05 or less, following correction for multiple comparisons, were deemed of interest and statistically significant.

### ELISAs

Concentrations of mouse IL-17A (Biolegend, #432504), IFNγ (Biolegend, #430804), IL-2 (Biolegend, #431004) and IL-22 (Biolegend, #436304), and of human IL-17A (Invitrogen #BMS2017) were determined in cell culture supernatants by ELISA, as per the manufacturer’s guidelines.

### IMMUNOHISTOCHEMISTRY

Lymph nodes were fixed in 10% neutral buffered formalin and embedded in paraffin. 10 μm sections were deparaffinized and antigens retrieved by microwaving in tri-sodium citrate buffer, pH6 for ten minutes, or by ten minutes’ incubation with 5mg/ml proteinase K (ThermoFisher UK #AM2548). Slides were blocked (Avidin/Biotin Blocking Kit, Vector Laboratories, #SP-2001) and incubated overnight with rabbit anti-mCRAMP (1/250; Innovagen, #PA-CRPL-100). Slides were then incubated with an anti-rabbit HRP-conjugated secondary antibody (1/500; Dako, #P0217) or AF488-conjugated antibody (1:500, ThermoFisher UK #A11034). Some sections were developed with diaminobenzidine (ImmPACT DAB Peroxidase (HRP) Substrate, Vector Laboratories, #SK-4105) and others were also incubated with an AF647 anti-neutrophil antibody (1:50, Biolegend UK #127609). Slides were scanned on a ZEISS Axio Scan.Z1 slide scanner.

### HEAT-KILLED SALMONELLA MODEL

Heat-killed salmonella was a kind gift from Professor Andrew Macdonald, MCCIR, University of Manchester. 12.5 μg heat-killed *Salmonella typhimurium* in 50 μl PBS was injected subcutaneously in the top of each hind paw. Mice were monitored and the draining popliteal lymph nodes removed after 1, 3 or 7 days. Nodes were either fixed in 10% neutral buffered formalin and embedded in paraffin for sectioning, or single cell suspensions were prepared as described previously for flow cytometric analysis.

### STATISTICS

All data shown are expressed as individual data points with line at median. Analysis was performed with GraphPad Prism software. Multiple groups were compared by one- or two-way analysis of variance tests with either Bonferroni or Dunnett post-tests. Two groups were compared with paired Student’s *t*-tests. A minimum of three mice was used for *in vitro* experiments, in individually performed experiments. Details of sample sizes are included in all figure legends.

## ACKNOWLEDGEMENTS

We thank the QMRI flow cytometry facility for help and advice, and Dr Robert Gray and Professor Julia Dorin, both University of Edinburgh, for helpful discussions.

## AUTHOR CONTRIBUTIONS

Conceptualisation: DM, DJD, EGF; Funding acquisition: EGF, DJD; Investigation: DM, KJS, VA, GH, LJJ, EGF; Methodology: AMcD, EGF; Project administration: EGF; Resources: AMcD, DJD, EGF; Supervision: DJD, EGF; Writing – original draft: DM, EGF; Writing – review and editing: DM, EGF, DJD

